# Cryo-EM structure of amyloid fibril formed by α-synuclein hereditary A53E mutation

**DOI:** 10.1101/2022.03.11.483992

**Authors:** Chuanqi Sun, Kang Zhou, Peter DePaola, Woo Shik Shin, Trae Hillyer, Michael R. Sawaya, Z. Hong Zhou, Lin Jiang

**Affiliations:** Department of Neurology, David Geffen School of Medicine, UCLA, Los Angeles, CA 90095, USA; California Nano Systems Institute, UCLA, Los Angeles, CA 90095, USA; Departments of Biological Chemistry and Chemistry and Biochemistry, Howard Hughes Medical Institute, UCLA-DOE Institute, UCLA, Los Angeles, CA 90095, USA; Department of Pharmaceutical Sciences, College of Pharmacy, Northeast Ohio Medical University, Rootstown, OH 44272, USA

## Abstract

Synucleinopathies, including Parkinson’s disease (PD), dementia with Lewy bodies (DLB), and multiple systems atrophy (MSA) have the same hallmark pathologic feature of misfolded α-synuclein protein accumulation in the brain. PD patients who carry α-syn hereditary mutations tend to have an earlier onset and more severe clinical symptoms and pathology than sporadic PD patients who carry wild-type (WT) α-syn. Therefore, revealing the structural effect of α-syn hereditary mutations on the wild-type fibril structure can help us understand synucleinopathies’ structural basis. Here, we present a 3.38 Å cryo-electron microscopy structure of α-synuclein fibrils containing the hereditary A53E mutation. The A53E fibril is symmetrically composed of two protofilaments, as are many other synucleopathic structures – including WT. Interestingly, the interface between the protofilaments in A53E has significantly less buried surface area than all other documented fibril structures of α-syn and its other mutants. The A53E fibril also exhibits slower formation/growth in *in vitro* fibrillation experiment compared to other mutants. This implies that the structural differences - both in the protofilament and between each protofilament of A53E – change the aggregation mechanism, or in the least, its kinetics of formation. These differences influence the molecular characteristics of each fibril mutant and likely plays a macro-scale role in progressing one clinical pathology over another.

## Introduction

Synucleinopathies, including Parkinson’s disease (PD), dementia with Lewy bodies (DLB), and multiple systems atrophy (MSA), are all linked to the accumulation, deposition and dysfunction of misfolded alpha synuclein(Spillantini, 1999; Spillantini et al., 1998). There is compelling evidence that the hallmark feature of Parkinson’s disease is the aggregation of alpha synuclein which makes up the major contents of Lewy bodies (LBs) and Lewy neurites (LNs). In the patients diagnosed with MSA, the same fibrillar form of alpha synuclein from Glial cytoplasmic inclusions (GCIs) is found in the oligodendrocytes of white matter tracts (Crowther et al., 2000; Spillantini & Goedert, 2000). Moreover, α-synuclein’s (SNCA) genetic variability (including single point mutation and genomic duplication or triplication of *SNCA*) is robustly linked to familial parkinsonism play crucial molecular roles in the early onset of PD(Fujioka et al., 2014; Goedert, 1997; Nussbaum & Polymeropoulos, 1997; Pankratz et al., 2009; Polymeropoulos et al., 1997). So far, multiple single point mutations have been found to cause familial Parkinson’s disease, including A30P, E46K, H50Q, G51D, A53E and A53T, leading to an earlier onset and more severe clinical symptoms and pathology (Appel-Cresswell et al., 2013; Hoffman-Zacharska et al., 2013; Kruger et al., 1998; Lesage et al., 2013; Pasanen et al., 2014; Pettersen et al., 2004; Polymeropoulos et al., 1997; Zarranz et al., 2004).

α-synuclein A53E is a novel hereditary mutation and was first discovered from a biopsy of a Finnish patient with atypical Parkinson’s disease (PD)(Pasanen et al., 2014). Previous studies have shown that the A53E mutation delays α-synuclein fibril formation and enhances toxicity in the cell cultures under cellular stress conditions(Mohite, Navalkar, et al., 2018). *In vitro* studies indicate that A53E mutation have a lower membrane binding affinity than the wild-type protein (Ghosh et al., 2014). Moreover, the patient with this novel *SNCA* A53E mutation showed highly abundant α-synuclein pathology throughout the brain and spinal cord with both MSA and PD features. Different PD mutations have been linked to different forms of disease with variable pathologies. The A30P and H50Q mutations are linked to patients with classic PD, while E46K is linked to DLB, and A53E, A53T, and G51D, bring about MSA and severe PD symptoms (Guan et al., 2020; Martikainen et al., 2015) (Mohite, Kumar, et al., 2018; Schneider & Alcalay, 2017). This noticeable variation in severity and disease development between different a-syn mutations highlights the importance of key residues that influence the fibril fold and overall negative effects caused by fibrils. By characterizing and determining the near-atomic resolution structure of fibrils with mutated a-syn, we may help to understand the molecular mechanism in the formation and spreading of synucleinopathy.

Our previous work determined cryo-EM structures of the full-length α-synuclein fibrils, which showed two distinct polymorphs, termed: rod and twister(Li et al., 2018). The rod protofilaments contain ordered residues 38-97 and residues 50-57 from the preNAC region form the interface, whereas the twister polymorph contain ordered residues 43-83 and residues 66-78 from the NACore form the interface. From the wild-type structure we know A53 is located at the hydrophobic interface of protofilament. This indicates that A53E mutation may alter key contacts at the protofilament interface, potentially leading to a different fibril structure and pathology. Therefore, we planned to determine the cryo-EM structure of α-syn fibrils containing the A53E hereditary mutation. Using the wild type and other mutant fibril structures, we can comparatively show the structural similarities and differences in an effort to provide insight into the key interactions between residues within fibrils, their influence on their fibrillar energetic fold, and speculate how single mutations at key locations within a-synuclein correlates with the observed severity between wild-type and heritable Parkinson’s.

## Results

### Cryo-EM structure and architecture of A53E fibril

In order to further study the influence of this new hereditary mutation on the α-syn amyloid fibril formation, we expressed and purified recombinant full-length α-syn containing A53E hereditary mutation and grew fibrils using our previously established protocol (Li et al., 2018). 2D Classification of the cryo-EM images revealed that the fibril sample is morphologically homogeneous with one major specie (Figure 1-figure supplement 1). After helical reconstruction of the major specie with Relion 3.1, we obtained the cryo-EM structure of A53E α-syn fibrils to a resolution of 3.38 Å. The A53E fibril polymorph has a pitch of ~880 Å with a width of ~12 nm, the fibril contains two intertwined protofilaments related by an approximate 2_1_ screw axis of the symmetry with a helical rise 2.4 Å and the helical twist between α-syn subunits is 179.5° (Figure 1B, C, Figure1-figure supplement 2A, B).

**Figure 1.**
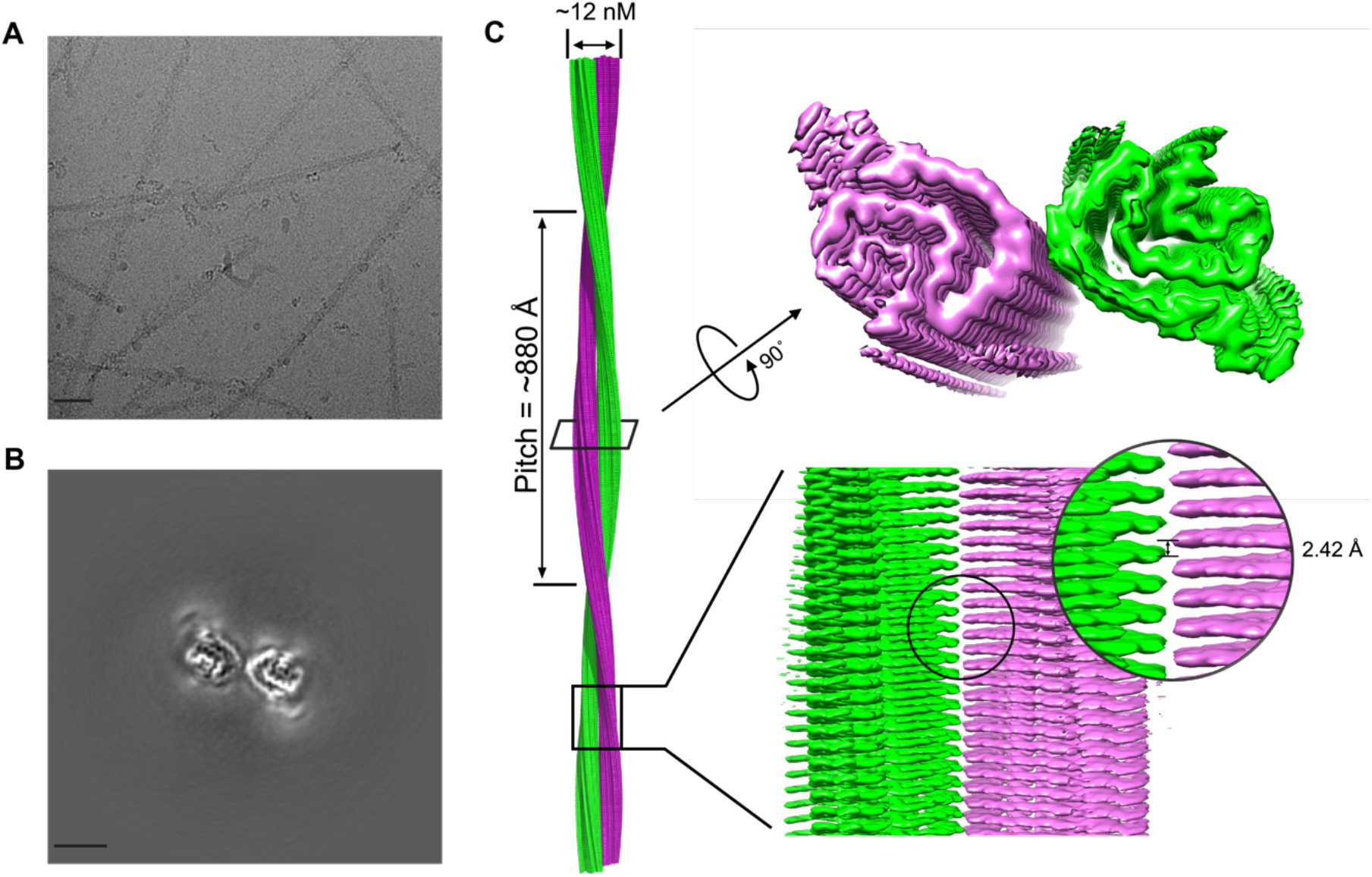
Cryo-EM structure of A53E polymorphic fibril. (**A**) A raw cryo-EM image of amyloid fibrils from full-length human A53E a-syn. Scale bar, 500 Å. (**B**) Cross-sectional view of the 3D map of these fibrils showing two protofibrils forming a dimer. Scale bar, 500 Å. (**C**) Cryo-EM 3D reconstruction density map of the A53E α-syn fibril. Fibril width, length of half pitch (180° helical turn), helical rise is indicated. The two protofilaments are colored in green and purple, respectively.

Based on the high-quality cryo-EM density map we obtained, we unambiguously built an atomic structure model for the A53E fibril (Figure 2A). The atomic model reveals that the A53E fibrils are wound from two protofilaments, 62 residues out of a total of 140 amino acids of α-syn from Leu38 to Gln99 and form a Greek-key-like fibril core, which is similar to that of a WT α-syn fibril structure with two turns. Each α-syn molecule comprises seven in-register parallel β-strands (i.e., residues (i.e., residues 48-50 (β1), 52-56 (β2), 61-64 (β3), 69-72 (β4), 75-77 (β5), 81-83 (β6), 94-96 (β7)). The A53E fibril core features a loosely fold contains 7 short β-strands, which is different from previous wild-type α-syn fibril core contains a long β-strand(β1) and 4 short β-strands. A53E fibril core is stacked every 4.8 Å along the fibril axis on a two-dimensional layer, forming mature β-sheet fibrils up to hundreds of nanometers (Figure 2C, D, E). Packing schema show that the stabilized A53E fibril has a hydrophobic core surrounded by hydrophilic residues (Figure 2B). Data collection and refinement statistics are summarized in Table 1.

**Figure 2.**
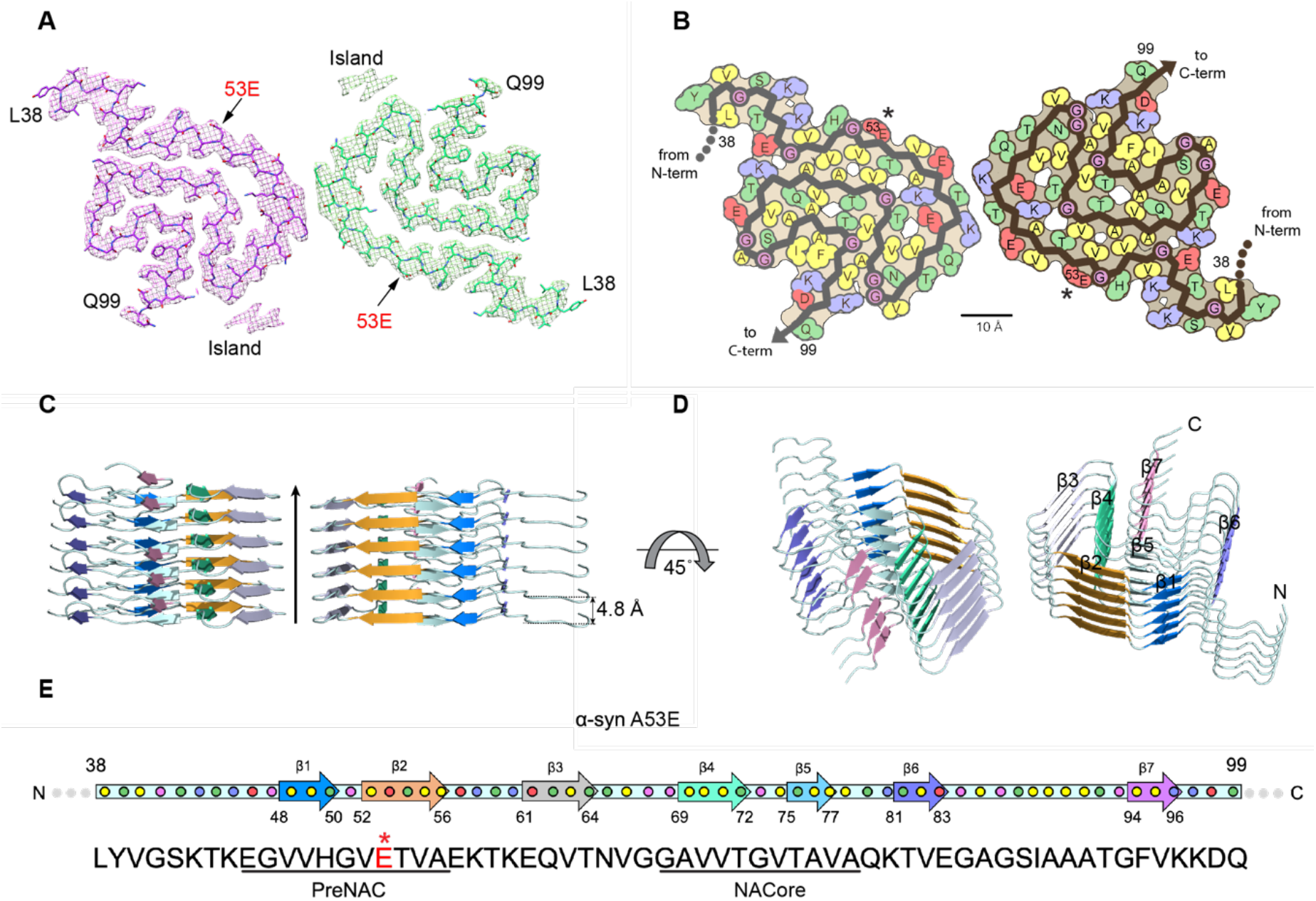
Overall structure of the α-syn A53E fibril. **(A)** Top view of the A53E fibril. One layer of the structure is shown, which consists of two α-syn molecules covering residues 38–99. The two molecules are colored differently. **(B)** Schematic representation of fibril structure with amino acid side chains colored as follows: hydrophobic (yellow), negatively charged (red), positively charged (blue), polar, uncharged (green). **(C)** Side views the multi-layer structure of A53E fibril, the distance between neighboring layer in one protofilament is indicated. **(D)** Ribbon representation of the structure of an A53E fibril core in b), β-strands on the top layer are shown. **(E)** Primary and secondary structure of a-syn A53E fibril. Arrows in the secondary structure indicate that conservative residues adopting β-strand conformation, and each colored dot represents a residue. PreNAC comprises residues 46–56; NACore comprises residues 68–78. The A53E mutation lies in the preNAC region, asterisk indicate the location of residue 53 (red).

**Table 1.** Cryo EM data collection, refinement and validation statistics.

| **A53E fibril** |  |
| --- | --- |
| **Data collection** |  |
| EM equipment | Titan Krios |
| Voltage (KV) | 300 |
| Detector | K2 summit |
| Nominal Magnification | ×130,000 |
| Pixel size (Å) | 1.07 |
| Total electron dose (e<sup>-</sup>/Å<sup>2</sup>) | 47 |
| Frame exposure time (s) | 0.2 |
| No. of movie frames | 35 |
| Defocus range (μm) | -1.5 ~ -3.0 |
| **Reconstruction** |  |
| No. of micrographs collected | 3,289 |
| No. of micrographs used | 2,767 |
| No. of filaments picked | 46,626 |
| No. of segments for 3D auto-refine | 31,916 |
| Resolution (Å) | 3.38 |
| Map sharpening B-factors (Å<sup>2</sup>) | -125 |
| Symmetry for final map | C1 |
| Helical rise (Å) | 2.42 |
| Helical twist (°) | 179.5 |
| **Atomic model** |  |
| No. of atoms | 2,586 |
| No. of protein residues | 372 |
| R.m.s deviations |  |
| Bonds length (Å) | 0.003 |
| Bonds angle (°) | 0.471 |
| Ramachandran plot statistics (%) |  |
| Preferred | 85.56 |
| Allowed | 14.44 |
| Outlier | 0 |
| Rotamers outliers (%) | 0 |
| C-beta deviations | 0 |
| Bad bonds (%) | 0 |
| Bad angles (%) | 0 |
| MolProbity score | 2.29 |
| Clash score | 13.48 |
| PDB code | 7UAK |
| EMDB code | EMD-26427 |

Additionally, we observed two unidentified densities flanking the two protofilaments in the A53E fibril, which we term ‘islands’. The side chain located on the island, possibly from residues 106-109 or 111-114, could form a hydrophobic interaction with residues T^64^-N^65^-V^66^ on the A53E beta-arch to stabilize A53E fibrils. The similar islands are also observed in the structure of α-synuclein hereditary disease mutant H50Q we determined. The presence of islands in different fibrils also reflects the structural diversity of amyloid protein in addition to the fibril conserved kernel and may also be related to the significant differences in α-syn pathology (Figure 2-figure supplement 2A, B).

### Comparison of A53E and wild type α-Syn protofilament folds

Overall, the hereditary mutation A53E compared with wild-type rod and twister fibril polymorphs, has a common conserved β-arch-like protofilament kernel structure, but different inter-protofilament interface (Figure 3D). In the WT rod structure, A53E is involved in the steric-zipper interface of the two protofilaments formed by preNAC region residues 50-57 (^50^HGVA/ETVAE^57^), which is assumed to be disrupted by A53E mutation. In the WT twister polymorph, NACore residues 66-78 (^66^VGGAVVTGVTAVA^78^) form a homotopical steric-zipper interface to bundle the two protofilaments together. However, the A53E fibril contains a protofilament interface formed by only two residues ^59^TK^60^ not previously seen in the wild-type α-Syn structures. Residues ^59^TK^60^ are located at a sharp turn in both protofilaments, and obviously, the interface between protofilaments is extremely small and no has obvious interaction (Figure 3 A, B, C). Further analysis revealed the buried surface area of be 10.3 Å^2^ and has a shape complementarity of 0.06. The A53E fibril interface is very small compared to the more extensive preNAC and NACore interfaces seen in our previous wild-type structures. Even compared with the Ac-A53T protofilament interface, which contains the same two residues interfaced, A53E still has the smallest fibril protofilament interface observed to date (Figure 3-Figure supplement 3 A, B, C, D).

**Figure 3.**
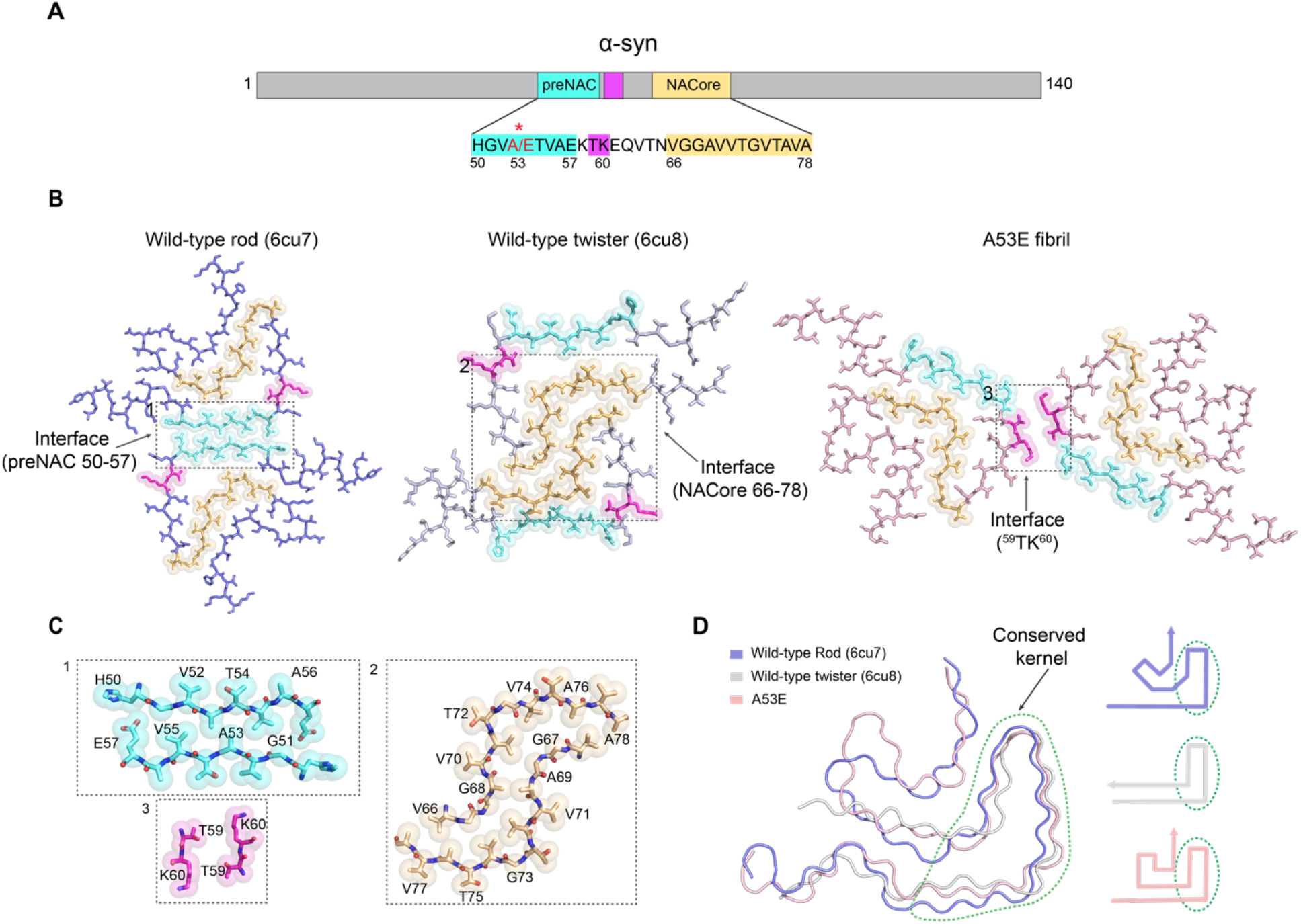
Comparison Wild type and A53E polymorphs. **(A)** Primary structure schematic highlighting residues of the conserved kernel (50–78) that form protofilament interfaces in α-syn polymorphs. The preNac region is shown in cyan, residues ^59^TK^60^ are shown in purple, and NAcore region is shown in orange. **(B)** Protofilamental interface of the WT rod, twister and A53E fibrils. Residues involved in the interfaces are highlighted with spheres. The fibril interface of WT rod fibril is colored in cyan; that of the WT twister fibril is in magenta; that of the A53E fibril is in orange. The interface and electrostatic interactions are zoomed in**(C)**. **(C)** Zoom-in views of the fibril interface interactions. Interface residues are labeled. d), Compare structures of every single α-syn subunit from the A53E, wild type rod, and wild type twister fibrils. A53E fibril is in lightpink; Wild type rod fibril is in slate. E46K fibril is in gray. The conserved kernel region with a similar structure shared by three different fibril is marked with a green dashed circle.

The second key difference between A53E and wild-type structures is their pattern of electrostatic interactions. In the wild-type rod fibril structure, residues E46 and K80 form a salt bridge, which is crucial for the stabilization of the Greek-key-like structure conformation (Figure 4 B, E). In the wild-type twister fibril, the E57-K58 also form an essential surface-exposed salt bridge to stabilize fibril conformation (Figure 4 C, F). In contrast, in the A53E fibril, the A53E mutation rearranges the electrostatic interactions in the fibril and trigger obvious changes in the electrostatic interactions, only can see one new salt bridge formed by K96 and D98 residues, to stabilize the conformation of the C-terminal region (Figure 4 A, D).

**Figure 4.**
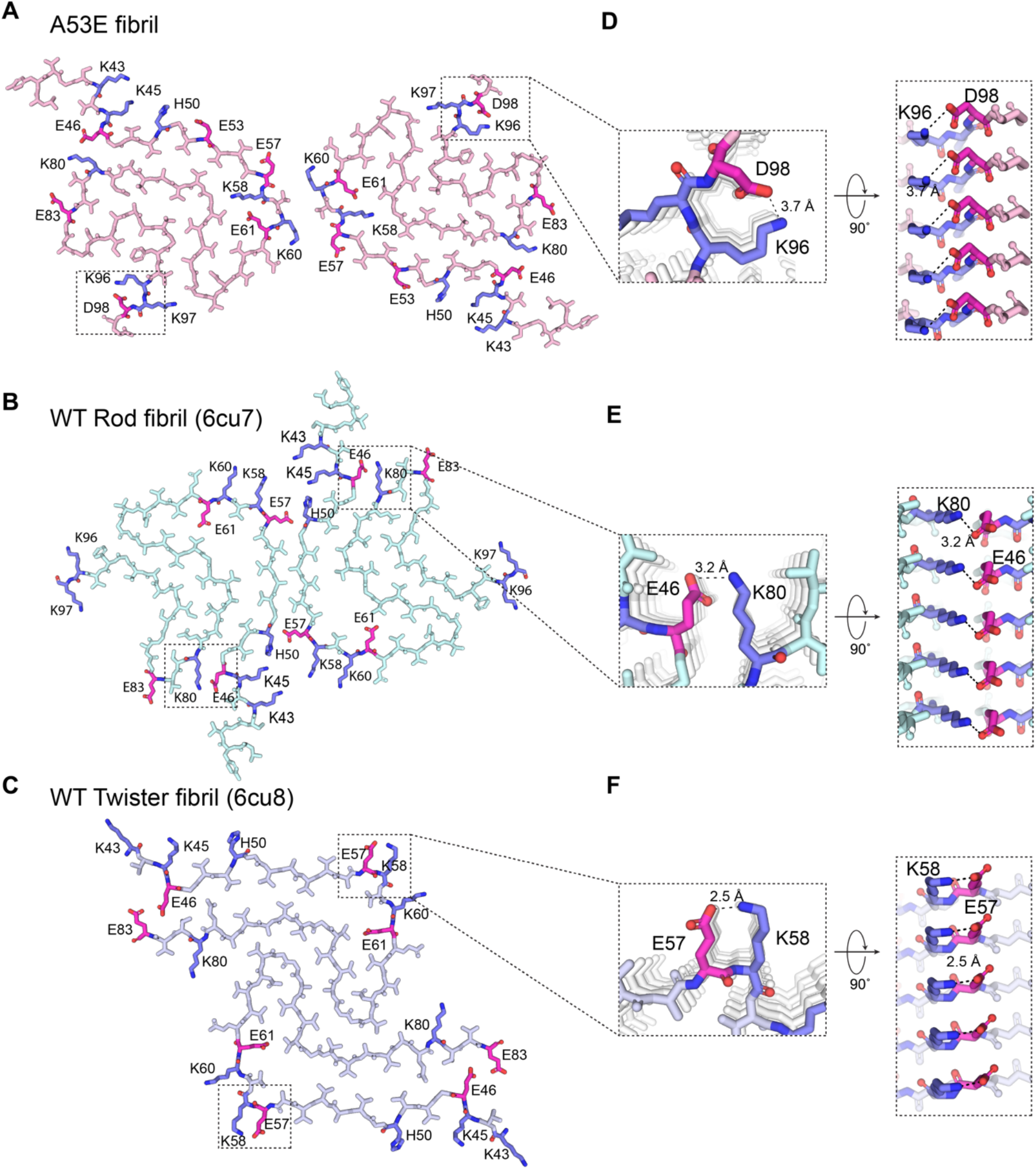
Electrostatic interactions in wild type and A53E α-syn fibril polymorphs. **(A-C)** One layer of A53E, wild type rod and twister fibrils are shown by sticks (colored by lightpink(a), palecyan(b) and bluewhite(c), respectively), charged and ionizable residues are colored differently, K and H are colored in slate, E and D are colored in magentas. d-f), A magnified top view of electrostatic interactions in A53E and wild type fibrils, where two pairs of amino acids (Lys 96 and Glu 98 in A53E; Glu 46 and Lys 80 in Rod; Glu 57 and Lys 58 in twister fibril) from opposing subunits form one salt bridges. A side view (right) highlighting a strong salt bridge between two residues from its opposing subunit, with a distance of 3.7, 3.2 and 2.5 Å (black).

### Cavity difference in α-syn fibril structures

The formation of cavity is a common feature of the α-syn fibril structures that have so far been determined. In our previously obtained wild type α-syn fibril structure, the cavity is surrounded by residues T54, A56, T59, E61, T72, G73, and T75 at the center of the β-arch. However, in the hereditary mutation A53E fibril, only residues T54, A56, K58, G73, V74 are involved in the formation of the cavity, making it smaller than the wild type one (Figure 2 A, B). Moreover, this is probably caused by conformation change of residues K58 and T59 side chain orientation. Compared with the wild-type and MSA brain extracted structure, the K58 residue side chain faces inwards towards the cavity, while T59 flips away from the cavity to the solvent. This inversion is speculated to promote the stability of the hereditary mutation fibril, and generate a more compact fibril core. Similarly, the orientations of side chains of K58 and T59 are flipped can also be seen in other mutants, such as H50Q, A53T fibril structures (Figure 2-Figure supplement 2 A-F) (Boyer et al., 2019; Sun et al., 2020).

In the cavity of A53E structure, we see an ambiguous density that is smaller than a similar ambiguous density found in our previous H50Q fibril structure. Moreover, given the resolution of A53E density map, we cannot determine whether this density is part of the protein structure or bound solvent molecules (Figure 1 C right-up). It can be inferred that the additional density between A53E and H50Q structures is distinct as they don’t align. The presence of a cavity in multiple α-syn fibril structures, as well as noticeable density inside their cavities might point towards the dynamics and conformational flexibility of the cavity near fibril core that triggers or contributes to the transition from monomeric to pathological a-syn.

### Comparison of the stability and pathological characteristics of WT and A53E α-syn fibrils

To access the role that the A53E hereditary mutation plays in fibril aggregation and stability, we first studied the kinetic propensity of A53E to aggregate compared to the wild type of protein. Our results indicate that, both A53E and WT aggregation proceeded with a typical sigmoidal curve. A53E reached maximum intensity in about 52 h, while it only took about 38 h for the wild type α-syn to reach maximum intensity. This shows that the aggregation rate of A53E is significantly slower than that of wild-type protein, consistent with previous other studies (Figure 5A). We next asked if the structural differences that we observe in the A53E versus wild-type fibrils affect their stabilities. We performed a sodium dodecyl sulfate (SDS) denaturation assay where both wild-type and A53E fibrils were incubated with various concentrations of SDS at 37 °C followed by thioflavin-T (ThT) fluorescence measurements. Our results demonstrate that A53E fibril is slightly more susceptible to chemical denaturation than wild-type fibrils (Figure 5B). Further solvation energy indicates that the free energy of A53E and wild-type fibrils has no significant difference, and that the A53E fibril energy can maintain fibril stability like wild-type one (Figure 5-figure supplement 1A-C, table 2). We speculate that the stability difference of A53E compare with wild-type fibril in the SDS denaturation is due to the A53E weak interface, whose meager interaction interface will show poor stability in the presence of strong denaturants.

**Figure 5.**
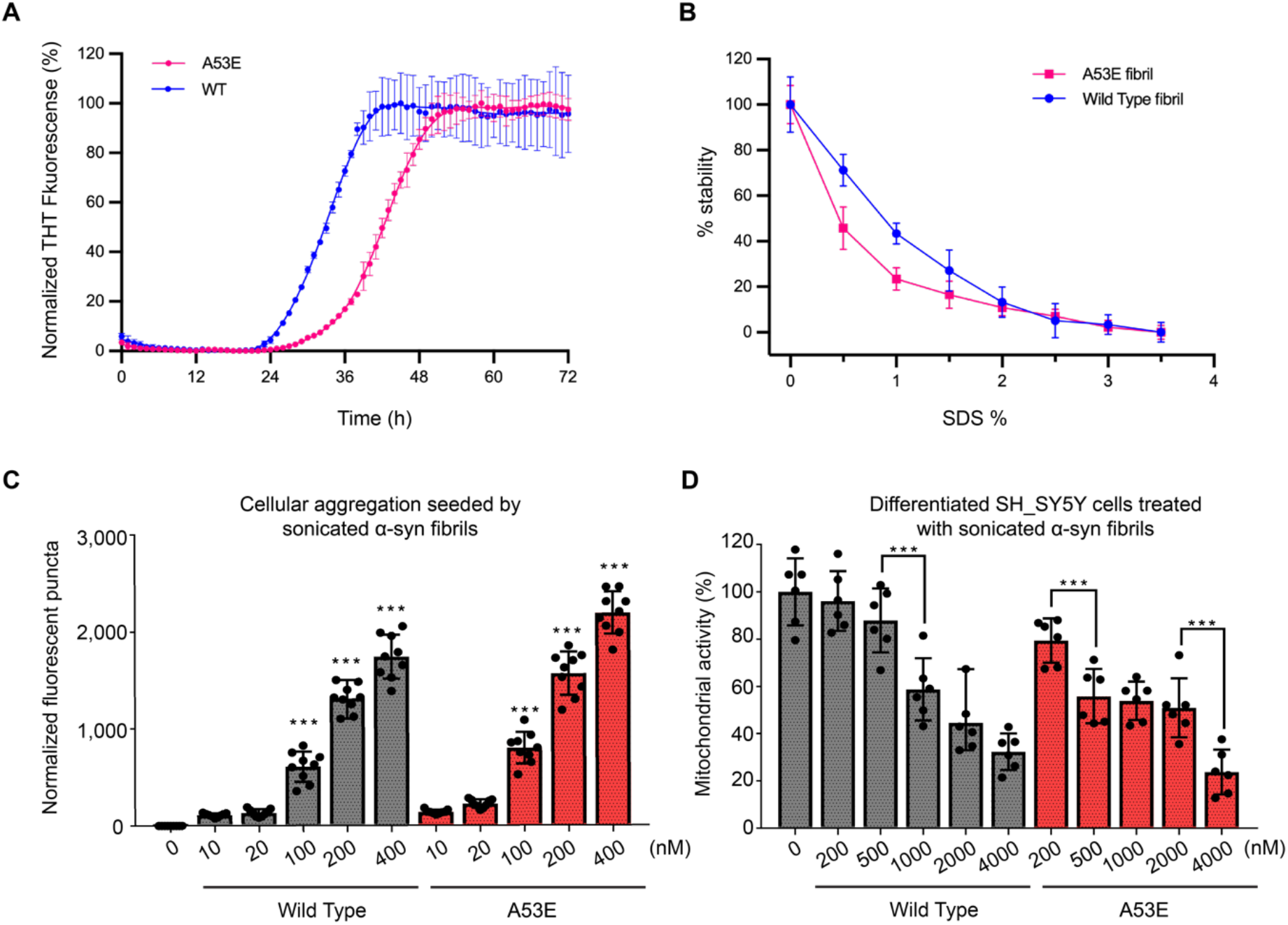
Comparison of the stability and pathological characteristics of WT and A53E α-syn fibrils. **(A)** THT assay measuring kinetics of A53E and wild-type α-syn aggregation. Wild-type aggregation plateaus at 38 h whereas A53E aggregates slower and plateaus at 50 h. Data are shown as mean ± s.d., n = 3 independent experiments. **(B)** Stability assay of A53E and wild-type α-syn. A53E and wild-type fibrils were heated to 37 °C and incubated with varying concentrations of SDS. A53E fibrils are more unstability to SDS than wild-type fibrils. Individual triplicate measurements are shown, and the plotted line represents the average of the triplicates. Data are shown as mean ± s.d., n = 3 independent experiments. **(C)** HEK293T α-syn YFP biosensor cells were used to perform the cell seeding assay of wild-type α-syn and A53E fibril. Each of sonicated fibril samples were transfected into biosensor cells using Lipofectamine 3000. After 48 h, the fluorescence microscopy images of α-syn biosensor cells were taken and the number of fluorescent puncta indicating aggregated endogenous α-syn YFP was counted. The scale bar denotes 100μm. A53E fibrils have a higher seeding capacity than wild-type α-syn fibrils. Data are presented as mean ± SD (n = 9, ***p < 0.0005, one way ANOVA). **(D)** MTT assay were performed to measure the toxicity of wild-type and A53E α-syn in differentiated SH-SY5Y neuroblastoma cells. Each of mild sonicated wild-type and A53E α-syn fibril samples was added to the cell culture medium and after 72 h incubation, cell mitochondrial activity was measured via MTT assay. Smaller amounts of A53E fibrils significantly disrupt mitochondrial activity and are more toxic than wild type.Data are presented as mean ± SD (n = 6, ***p < 0.0005, one-way ANOVA).

**Table 2.** Comparative different a-synuclein fibril solvation energy calculations

| **Fibril structure** | **Ordered residues** | **Resolution(Å)** | **Energy/chain**<br>**(kcal mol<sup>-1</sup>)** | **Energy/residue**<br>**(kcal mol<sup>-1</sup>)** |
| --- | --- | --- | --- | --- |
| **A53E** | 62 (38-99) | 3.38 | -56.7 | -0.9 |
| **Wild-type Rod (6cu7)** | 60 (38-97) | 3.5 | -57.7 | -0.96 |
| **Wild-type Twist (6cu8)** | 41 (43-83) | 3.6 | -33.3 | -0.81 |
| **Ac-A53T (6lrq)** | 63 (37-99) | 3.5 | -56.5 | -0.9 |
| **H50Q wide fibril (6pes)** | 64 (36-99) | 3.3 | -58 (pf A) | -0.90 (pf A) |
|  |  |  | -54 (pf B) | 1.00 (pf B) |
| **E46K (6ufr)** | 63 (36-98) | 2.5 | -66 | 1.06 |
| **(pf, protofilament)** |  |  |  |  |

With this information about its lower stability, we examined the cellular seeding and toxicity of A53E and wild-type fibrils using HEK293T α-syn-A53T-YFP biosensor cells and SH-SY5Y cells, respectively. The results indicated that A53E fibrils have higher seeding capacity in α-syn-A53T-YFP biosensor cells and significantly higher cytotoxicity in SH-SY5Y cells (Figure 5C, D, Figure5-figure supplement 2). These results are similar to previous studies demonstrating the potential pathogenic enhancement of A53E versus wild-type α-syn fibrils (Mohite, Kumar, et al., 2018; Picillo et al., 2018).

## Discussion

The structural polymorphism of fibrils is a common feature of pathological amyloids, such as α-syn, Tau, and Aβ. Various evidence has shown that the synuclein fibrils of synucleinopathies, which include MSA, Parkinson’s disease, Parkinson’s disease with dementia, and dementia with Lewy bodies (DLB) are distinct(Schweighauser et al., 2020). The diversity of fibril structure has also shown us that the polymorphism of fibrils can cause significant differences in biological activity and clinical pathology(Boyer et al., 2020; Healey et al., 2016; Shewmaker et al., 2011; Tycko, 2015)(Falcon et al., 2018; Mehra et al., 2021; Schweighauser et al., 2020). We hypothesize that the structural variation in fibril structures could lead to different disease pathologies, and thus plan to explore structures of mutant synuclein fibrils to elucidate the structure activity relationship of synuclein fibrils in toxicity and pathology. Supporting this, multiple near-atomic fibril structures have suggested that hereditary mutations may disrupt the stability of wild type fibril fold confirmation, such as H50Q, E46K, A53T, G51D, etc., which show distinct morphologies from wild-type fibrils(Aguirre et al., 2020; Boyer et al., 2020; Sun et al., 2020; Sun et al., 2021; Zhao et al., 2020). With our newly determined structure of A53E, we can further contribute to the characterization of each mutant fibril structure and infer what properties influence its pathology.

In this study, we determined the cryo-EM structure of α-syn fibril with hereditary mutation A53E. Structure alignments of A53E and wild-type protofilaments reveal that A53E adopt a similar conserved kernel β-arch-like fold. However, A53E forms a different and less stable protofilament interface compared to both wild-type rod and twister fibrils interface, which only utilizes two residues ^59^TK^60^ to assemble protofilaments. Moreover, the introduction of a negatively charged residue - glutamate - in the A53E structure tends to create a kinetics barrier that resulted in a significant decrease in the aggregation rate of α-syn fibril formation. The A53E mutation is commonly seen in mixed PD and MSA pathology in patients, as well as another α-syn mutant, G51D. Both mutants introduce negative charge at a residue near the same spatial region, which might influence the kinetic aggregation or stability of the fibril in such a way that predisposes the patient to these particular pathologies.

In our A53E mutant fibril structure, there is strong influence from the electrostatic interactions that maintains the stability of the protofilament. For example, E46 and K80 form a key salt bridge to stabilize the fibril core. But in A53E protofilament, those two residues side chain are too far away to form a strong salt bridge. In contrast, compare with wild-type rod fibril, a new salt bridge formed between K96 and D98 to stabilize the C-terminal region of A53E fibril. In contrast to these regions of strong interaction and stability, the A53E mutant also shares sites of relatively high flexibility. In every mutant fibril structure, there are positions where amino acid side chains flip opposite what is seen in the wild-type structures. The three sites where this occurs in A53E, and other mutants, are residues K58/T59, V74/T75, and T81/V82 (Figure 2-figure supplement 3). The first pair, K58&T59, is flipped from the outward & inward orientation to inward & outward orientation, which is similar to the H50Q and A53T protofilaments. We speculate that this inversion may cause the cavity seen in mutant fibrils to become smaller, which might accommodate fewer co-factors or adopt more rigid conformation, making the fibril structure more compact. The second pair of residues T81 & V82 also undergoes a similar inversion, where the hydrophobic side chain of V82 is flipped to the inward side, which increases the hydrophobic interaction of the fibril core. While these flips don’t entirely change the overall fold between each mutant, it does provide insight into the sequence positions where most strain occurs. In all three cases, the flipped sites occur at a turn in the layer and are places where we’d expect the most torsional strain when conforming to a flat layered structure. We guess that the minute change in a single amino acid causes a downstream shift in the overall structure, allowing the most strained amino acid conformations to relax into another spatial orientation compared to wild-type. By focusing on regions of variable instability between mutants with single point mutations, we can identify regions of key, flexible sites which may be of interest for those looking to design therapeutics to interact and destabilize a-syn fibrils.

In the first ex vivo α-syn fibril structure, which was extracted from the brains of patients with MSA by Schweighauser et al, the brain-derived Type I and Type II filaments have a cavity, that contains nonproteinaceous density(Schweighauser et al., 2020). Type I filaments have a larger cavity containing more additional densities than the cavity of Type II filaments. The authors also note that the density within the cavity could contain an unknown molecule that binds to the filaments. These finding indicate that a possible event of co-factor binding located near the cavity may contribute to maintaining fibril formation and/or stability. The side chain inversions which we have noted may also play a role in creating or closing the cofactor cavity by shifting neighboring residues to accommodate or exclude an accessory, or even competitive, cofactor. The binding of a cofactor molecule might nucleate the fold and pull a-syn residues around itself to transition into its filament conformation, a mechanism that begins with cofactor binding and forces the amino acids to pack around it, rather than an insertion of the cofactor into the fibril. The presence of a cavity in fibril structures may affect and contribute to the overall stability of the fibril and thus influence the symptoms and spread of fibril seeds within patients.

In summary, we determined the near-atomic structure of the α-syn family mutation A53E fibril. Compared with the wild-type fibril, it showed novel polymorphisms in different interface and electrostatic interactions. This structure enriches the fibril polymorphism of α-syn family mutants and provides a structural basis for us to further understand the influence of hereditary mutations on fibril formation and their pathogenic mechanisms.

## Acknowledgments

We thank Peng Ge and David Boyer for suggestions on this project. This work was supported by a grant from NIH (NIH AG 060149). We acknowledge the use of instruments at the Electron Imaging Center for Nanomachines (EICN) supported by UCLA and grants from the NIH (1S10OD018111 and 1U24GM116792) and the National Science Foundation (DBI-1338135 and DMR-1548924). A portion of this research was supported by NIH grant U24GM129547 and performed at the PNCC at OHSU and accessed through EMSL (grid.436923.9), a DOE Office of Science User Facility sponsored by the Office of Biological and Environmental Research.

## Author Contributions

L.J., C.S. and K.Z. designed experiments and performed data analysis. C.S. expressed and purified the α-syn protein. C.S. grew fibrils of α-syn and performed biochemical experiments. K.Z and Z.Z.H. performed cryo-EM data collection. C.S. and K.Z prepared cryo-EM samples,selected filaments from cryo-EM images,processed cryo-EM data and built the atomic models. W.S.S and T.H performed cell experiments. M.R.S. carried out solvation energy calculations. L.J. supervised and guided the project. C.S. wrote the manuscript with input from all authors.

## Materials and Methods

### α-syn expression and purification

Full-length α-syn wild-type and A53E mutant proteins were expressed and purified according to a published protocol ((Li et al., 2018)). Transformed bacteria were induced at an OD600 of ∼0.6-0.8 with 1 mM IPTG for 6 h at 30 °C. The bacteria were then lysed with a probe sonicator for 10 min in an iced water bath. After centrifugation, the soluble fraction was heated in boiling water for 10 min and then titrated with HCl to pH 4.5 to remove the pellet. After adjusting to neutral pH, the protein was dialyzed overnight against Q Column loading buffer (20 mM Tris-HCl, pH 8.0). On the next day, the protein was loaded onto a HiPrep Q 16/10 column and eluted using elution buffer (20 mM Tris-HCl, 1 M NaCl, pH 8.0). The eluent was concentrated using Amicon Ultra-15 centrifugal filters (10 NMWL; Millipore Sigma) to ∼5 mL The concentrated sample was further purified with size-exclusion chromatography through a HiPrep Sephacryl S-75 HR column in 20 mM Tris, pH 8.0. The purified protein was dialyzed against water, concentrated to 3 mg/mL, and stored at 4 °C. The concentration of the protein was determined using the Pierce BCA Protein Assay Kit (cat. No. 23225; Thermo Fisher Scientific).

### Fibril preparation and optimization

Both wild-type and A53E fibrils were grown under the same condition: 300 µM purified monomers, 15 mM tetrabutylphosphonium bromide, shaking at 37 °C for 2 wk.

### Negative stain transmission electron microscopy (TEM)

3.5 μl fibril sample was spotted onto a freshly glow-discharged carbon-coated electron microscopy grid for 2 min, then 5 μl uranyl acetate (2% in aqueous solution) was applied to the grid for 1 min. The excess stain was removed by a filter paper. Another 5 μl uranyl acetate was applied to the grid and immediately removed. The samples were imaged using an FEI T12 electron microscope.

### ThT kinetic assay

All the purified α-syn monomers (50 μM) were adequately mixed with 10 μM ThT and added into a 96-well-plate. Samples were incubated at 37 °C for over 3d with 600 r.p.m. double orbital shaking. The ThT signal was monitored using the FLUOstar Omega Microplate Reader (BMG Labtech) at an excitation wavelength of 440 nm and an emission wavelength of 490 nm.

### Measurement of α-syn fibril seeding using biosensor cells

HEK293T α-syn YFP biosensor cells were nicely provided from the laboratory of M. Diamond. The cells were grown in DMEM (Dulbecco’s modifications of eagle’s medium with L-glutamine & 4.5 g/l glucose) supplemented with fetal bovine serum (FBS), 100 units/ml of penicillin G, and 0.1 mg/ml of streptomycin sulfate, in a humidified atmosphere of a 5% CO_2_ at 37 °C. Trypsin-treated HEK293T cells were plated on collagen-coated flat 96-well plates at a density of 2.5×104 cells/well in 200 μl culture medium and incubated at 37 ºC in 5% CO_2_. α-syn fibrils were diluted with Opti-MEM (GIBCO) and sonicated for 10 min in an ultrasonic water bath. The α-syn fibril samples were then mixed with Lipofectamine 3000 (Thermo Fisher Scientific) and incubated for 15 min and then added to the cells. The actual volume of Lipofectamine 3000 was calculated based on the dose of 0.3 μl per well. α-syn fibrils aggregation in the biosensor cells was visualized at 48 hours by florescence microscopy and fluorescent images were processed in ImageJ to count the number of seeded cells. An Opti-MEM-treated control was used for normalization.

### MTT mitochondrial activity Assay

Human neuroblastoma SH-SY5Y cells were cultured in Iscove’s modified Dulbecco’s media supplemented with 15% fetal bovine serum (FBS), 1% L-glutamine, and 1% penicillin/streptomycin. For experiments, cells were transferred to collagen-coated flat 96-well plates at a density of 3.0×106 cells/well in 200 μl culture media and incubated at 37 ºC in 5% CO_2_ for 2 days. The cells were treated with different concentrations of wild-type and A53E α-syn fibrils (0.2 to 4 μM), which were sonicated in a water bath sonicator for 10 min before being added to the cells. After 72 h incubation, 10 μl of 3-(4,5-dimethylthiazol-2-yl)-2,5-diphenyltetrazolium bromide (MTT, 5 mg/ml) dye was added to each well, and the plates were incubated for 4 hours at 37 ºC in 5% CO_2_. After centrifugation at 1,500 rpm for 10 min, the supernatant was removed and 200 μl of dimethyl sulfoxide (DMSO) was added and the plates were gently shaken to solubilize the formed formazan for 30 min. The absorbance was measured using a microplate reader at wavelength 570 nm. The data were normalized to those from cells treated with Opti-MEM to obtain a value of 100%.

### Cryo-EM sample preparation, movies acquisition and drift correction

An aliquot of 2.5 μl of fibril solution was applied to a Quantifoil holey carbon grid (2/1, 300 mesh), that was glow discharged for 40 s with a PELCO Easy Glow system. The grid was blotted and plunge-frozen in liquid ethane with a Vitrobot IV (Thermo Fisher) at 4 °C under 100% humidity. The frozen grids were stored in liquid nitrogen before use. For data collection, the cryo-EM grids were loaded into an FEI Titan Krios electron microscope equipped with Gatan Quantum imaging filter (GIF) and a post-GIF K2 Summit direct electron detector. Movies were recorded as dose-fractionated frames in super-resolution mode by SerialEM automation software package(Mastronarde, 2005), with image shift induced-beam tilt correction. The slit width in the GIF system was set to 20 eV to remove inelastically scattered electrons. A total of 3,334 movies were recorded for the data set, the nominal magnification is ×130,000, corresponding to a calibrated pixel size of 0.535 Å on the specimen. An exposure time of 7 s was used at a rate of 0.2 s per frame, the dose rate per frame was set to 1.35 e−/ Å^2^, producing 35 frames and a total dosage of 48 e−/ Å^2^.

Frames in each movie were aligned for drift correction with the graphics processing unit (GPU)-accelerated program MotionCor2(Zheng et al., 2017). The first and last frame were discarded during drift correction. Two averaged micrographs, one with dose weighting and the other one without dose weighting, were generated for each movie after drift correction. The averaged micrographs were binned 2×2 to yield a pixel size of 1.07 Å. The micrographs without dose weighting were used for CTF estimation and particle picking, while those with dose weighting were used for particle extraction and in-depth processing.

### Cryo-EM data processing

The CTF estimation of each micrograph was performed by CTFFIND4 (ref.(Rohou & Grigorieff, 2015)). By discarding the micrographs either with underfocus values outside the allowed range (1.0–3.5 μm), or containing crystalline ice, we selected 2,767 good ones from the dataset. Then a total of 46,626 filaments were picked using crYOLO(Wagner et al., 2020; Wagner et al., 2019). Fibril particles were first extracted using a large box size (1,024 pixels) and a 10% inter-box distance in RELION3.1 (ref.(Scheres, 2012)), then the particles were subjected to two dimensional (2D) class averaging to determine the pitch. Helical parameters were deduced from the pitch with the assumption that each helix had a twisted twofold screw axis(Li et al., 2018). The 2D classes reveal that A53E α-syn forms fibrils of a single morphology with a pitch of ∼880 Å (Fig. S1c), which given the calculated helical twist of 179.5° (helical rise of 2.4 Å). Subsequently, we extracted all fibrils particles with a 512-pixel box size and 10% inter-box distance, yielding 226,604 particles. Using the calculated helical twist (179.5°) and helical rise (2.4 Å), these particles were subjected to a three-dimensional (3D) class averaging with a single class and a featureless cylinder created by EMAN2 (ref.(Tang et al., 2007)) as initial model. The cylinder was refined to a model in which two separated and twisted protofilaments could be seen. This model was then used to classify good and bad particles with a 3D class averaging with three classes. Particles in best class of the previous 3D classification were re-extracted with a 300-pixel box size for further 3D classification. Two additional 3D classifications were performed, and a final subset of 31,916 helical segments were selected and subjected to 3D auto-refinement, CTF refinement, and post-processing, yielding a map at 3.38 Å resolution (Fig. S2). Details of the data processing are summarized in Table 1.

The global resolution reported above is based on the ‘gold standard’ refinement procedures and the 0.143 Fourier shell correlation (FSC) criterion. Local resolution evaluation (Fig. S2) was performed with RELION3.1.

### Atomic Model Building

Atomic model building was accomplished in an iterative process involving Chimera (Pettersen et al., 2004), Coot (Emsley et al., 2010), and Phenix (Adams et al., 2010). Briefly, the structure of A53T α-syn (Sun et al., 2020) (PDB: 6LRQ) was fitted into the cryo-EM map as an initial model by Chimera. This fit revealed the mismatch segments of mainchain. After deleting the mismatch part, the model was refined by ‘real-space refinement’ in Phenix. We then manually built the missing residues and adjusted side chains to match the cryo-EM map with Coot. This process of real space refinement and manual adjustment steps was repeated iteratively until the peptide backbone and sidechain conformations were optimized. Ramachandran, secondary-structure restraints and NCS restraints were used during the refinement. Refinement statistics are summarized in Table 1. The model was also evaluated based on Ramachandran plots and MolProbity(Chen et al., 2010) scores.

## Supplementary Figures and Tables

**Figure 1-figure supplement 1.**
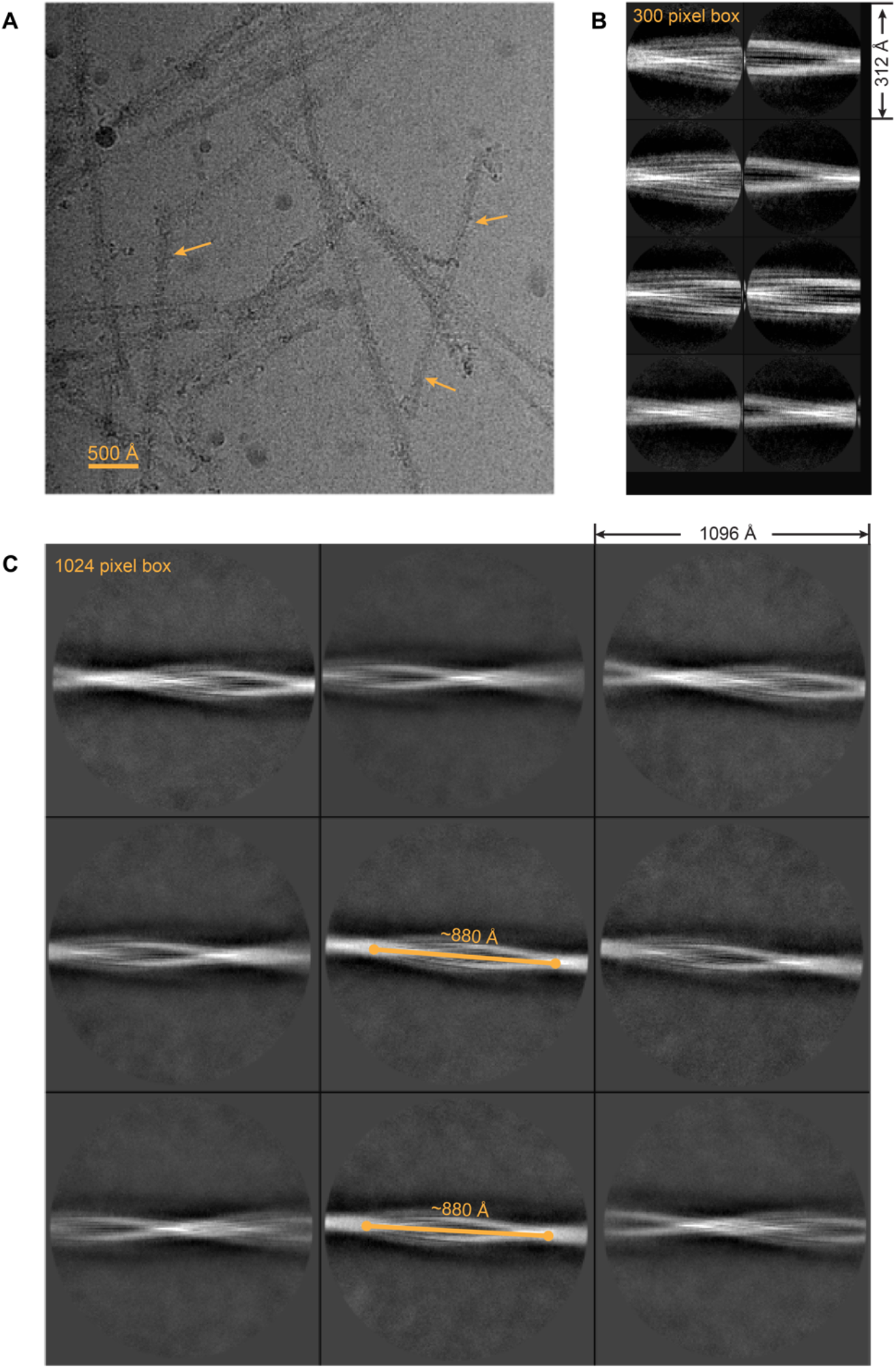
Cryo-EM image and 2D classification of A53E α-syn fibril. (**A**) A representative cryo-EM micrograph. Scale bar, 500 Å. (**B**) Representative 2D class averages at 300-pixel box (312 Å box length). (**C**), Representative 2D class averages at 1024-pixel box (1096 Å box length). Several classes show that the crossover distance of fibril is about 880 Å.

**Figure 1-figure supplement 2.**
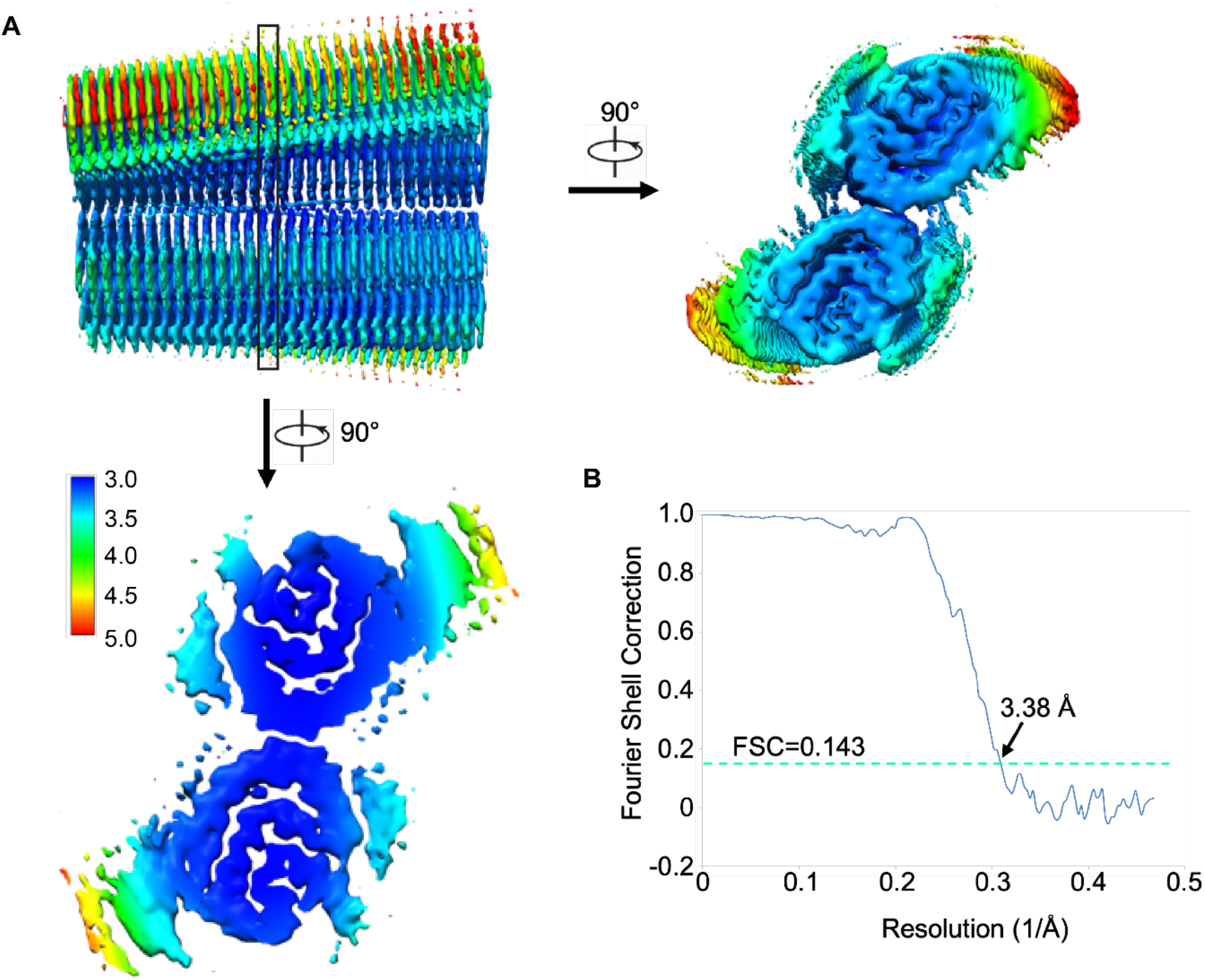
Resolution estimation of the cryo-EM map of A53E α-syn fibril. (**A**), Local resolution estimation. Density maps are colored based on the local resolution; the resolution of the core region is around 3.38 Å. The density map of A53E fibrils is colored according to local resolution estimated by ResMap. The enlarged cross sections and top view show the density map of two protofibrils. The color key on the left shows the local structural resolution in angstroms (Å) and the colored map indicates the local resolution ranging from 3.0 to 5.0 Å. (**B**), Gold-standard Fourier shell correlation curve. Gold-standard refinement was used for estimation of the density map resolution. The global resolution of 3.38 Å was calculated using a Fourier shell correlation (FSC) curve cut-off at 0.143.

**Figure 2-figure supplement 1.**
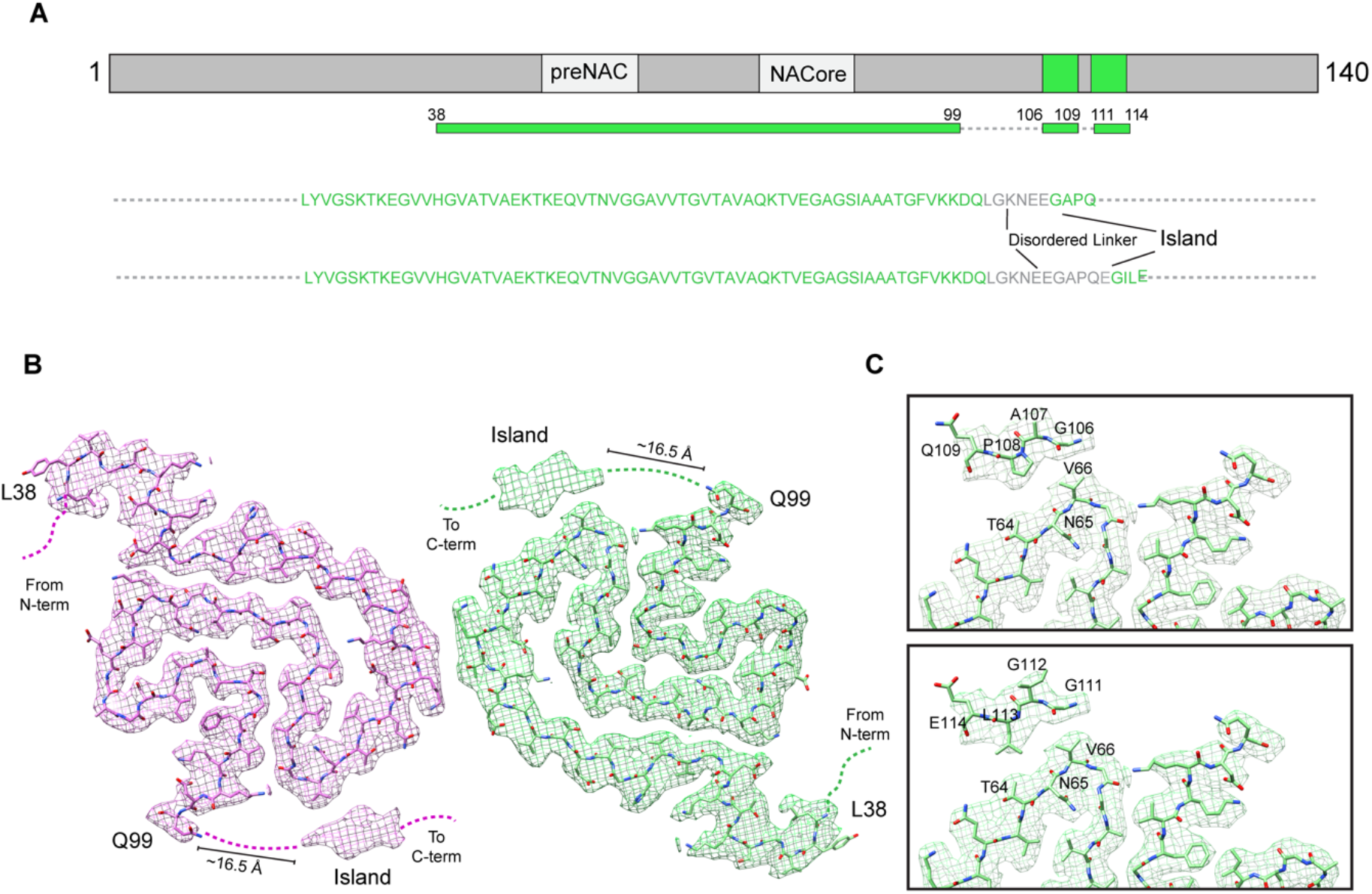
Speculative Atomic Model for Island suggested by the extra densities of A53E fibril. (**A**), Schematic diagram shows the core sequence of A53E fibril (^38^Leu - ^99^Gln) and the possible island sequence (^106^GAPQ^109^) and (^111^GILE^114^). (**B**), Illustration of possible regions from A53E protofilament fibril that could occupy Island. Some residues from C-terminal protofilament can give a reasonable explanation to the island. (**C**), Speculative model for Island in A53E protofiliment. Residues ^106^G^107^A^108^P^109^Q (up) or ^111^G^112^I^113^L^114^E (down) occupying the extra densities in our hypothetical model are labeled.

**Figure 2-figure supplement 2.**
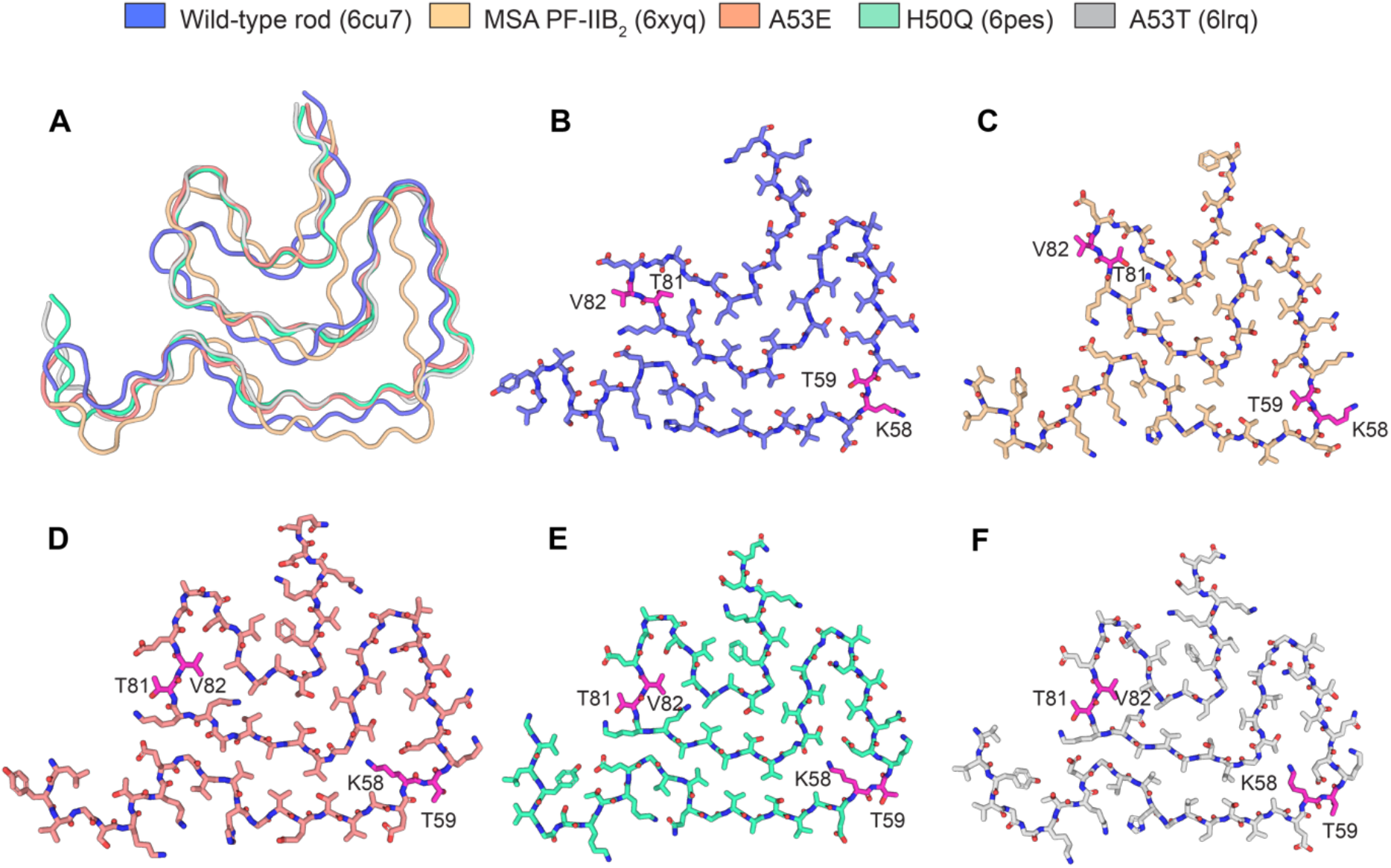
K58/T59 and T81/V82 show alternate conformations in different α-syn polymorphs. **A**), Structure alignment of wild type rod, MSA PF-IIB2, A53E, H50Q, and A53T polymorphs, fibrils are shown by Ribbon. **B-F**), Each fibril model is shown by sticks, residues K58/T59 and T81/V82 with conformation change are shown in red. Wild-type rod is shown in blue, MSA PF-IIB2 is shown in yellow, A53E is shown in pink, H50Q is shown in greencyan, A53T is shown in gray.

**Figure 2-figure supplement 3.**
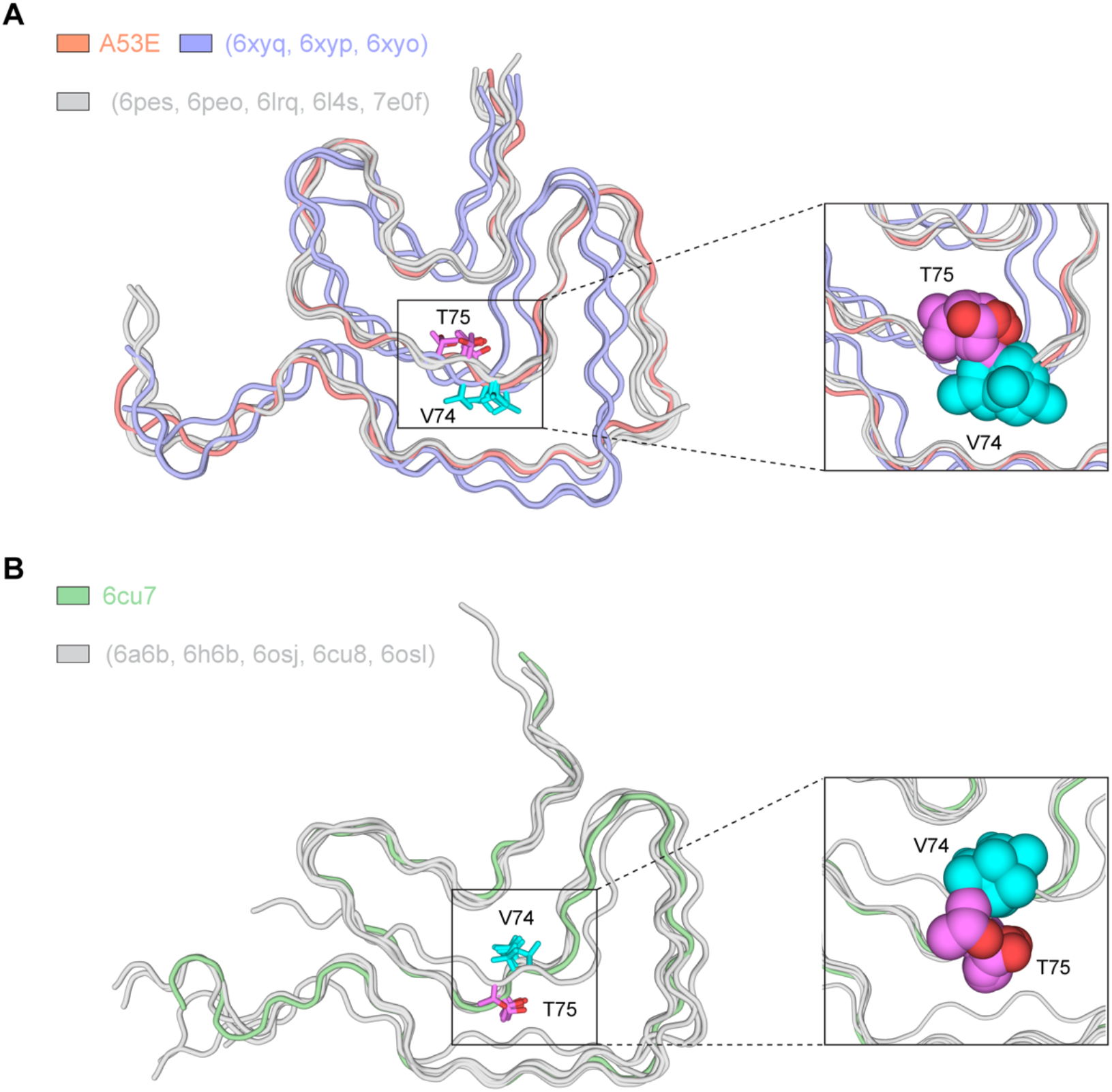
Structural alignment of different wild-type and mutant α-syn polymorphs. (**A**), Structural alignment of A53E protofilament (coral) with three types of patient extracted MSA filament (light blue) and hereditary disease mutant H50Q, E46K, A53T, G51D (gray) fibrils. Residues V74 (orchid) and T75 (cyan) with the same direction are shown by sticks, (right) zoom in two residues are labled. (**B**), structural alignment of full-length and truncated wild-type fibrils obtained from different growth conditions. Residues V74 (orchid) and T75 (cyan) are shown by sticks, (right) zoom in two residues are labled. Compared with all the mutant and MSA fibril, those two residues side-chain orientation is opposite.

**Figure 3-figure supplement 1.**
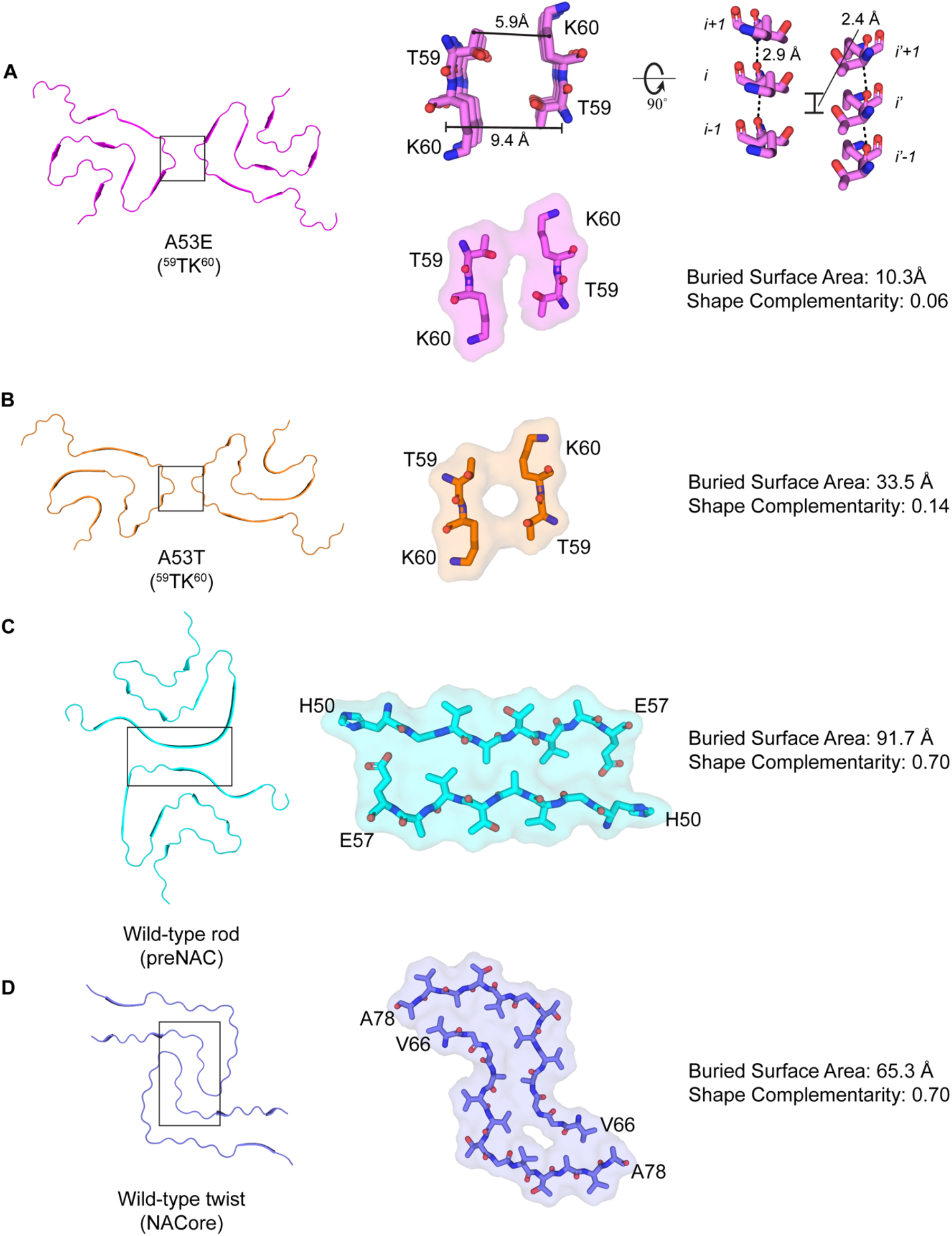
Comparison of protofilament interface of A53E, A53T, Wild Type rod and twist α-syn. (**A**), One layer A53E fibril overview(left), ^59^TK^60^ form the homointerface of the fibril and the distance between mated sheets of two protofilament is about 9.4 Å, the distance between Cγ atom of T59 and the corresponding K60 Cγ atom of the other layer is about 5.9 Å (right up). ^59^TK^60^ van der Waal’s surface, buried surface area and shape complementarityis shown (right down). (**B, C, D**), A53T, wild-type rod and twist one layer fibril overview(left), zoom-in views of A53T (^59^TK^60^), preNAC and NACore van der waal’s surface. Buried surface area and shape complementarity are shown, respectively.

**Figure 5-figure supplement 1.**
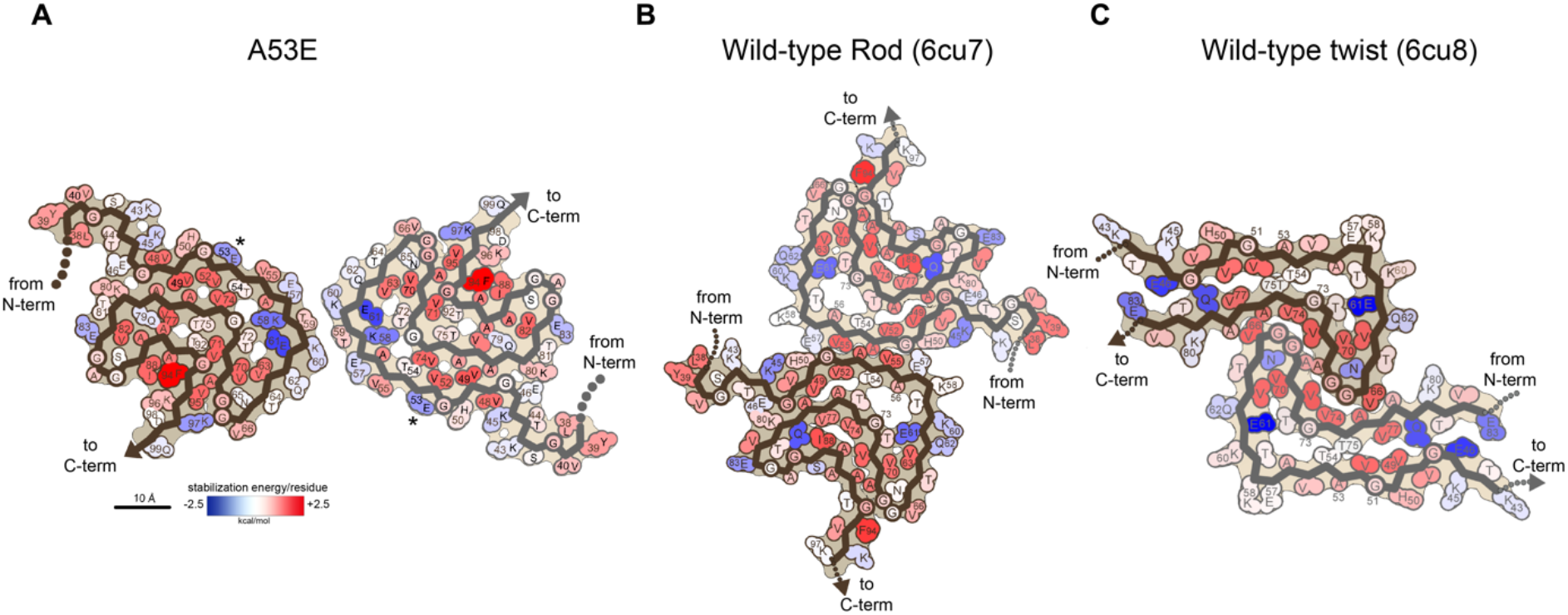
Schematic free energy of stabilization maps for the wild-type and A53E polymorphs. (**A-C**) Solvation energy maps of fibril structures. The stabilizing residues are red; the de stabilizing residues are blue.

**Figure5-figure supplement 2.**
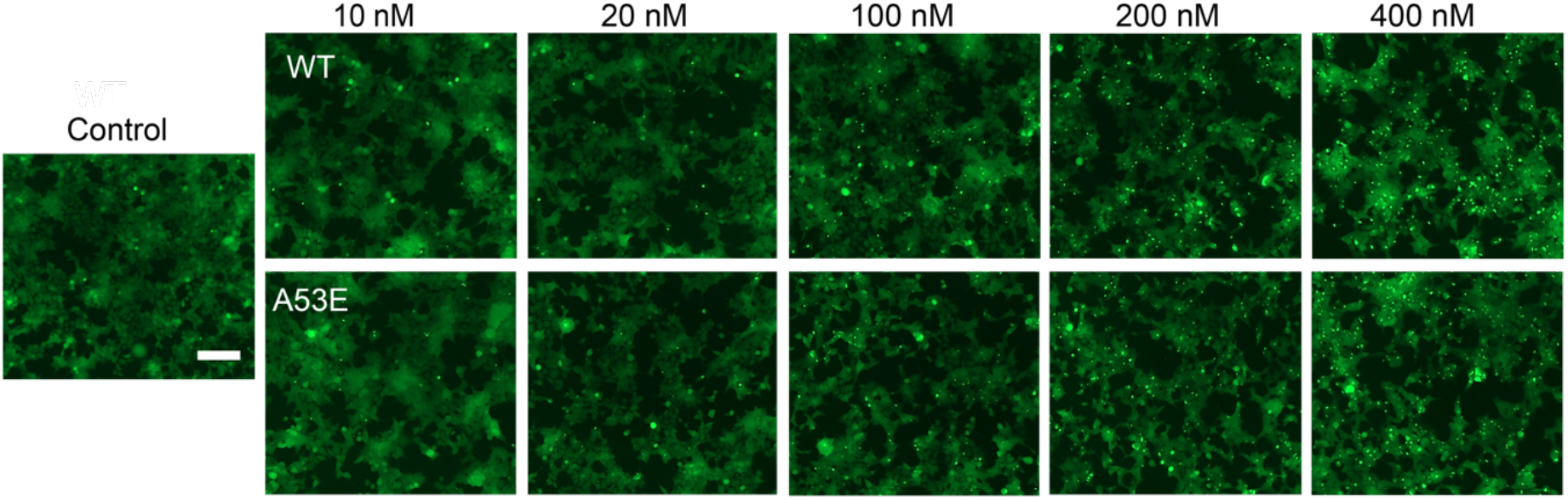
Fluorescence microscopy images of HEK 293T α-syn YFP biosensor cells were used to perform the cell seeding assay of wild-type α-syn and A53E fibril. Each of sonicated fibril samples were transfected into biosensor cells using Lipofectamine 3000. After 48 h, the fluorescence microscopy images of α-syn biosensor cells were taken. The scale bar denotes 100 μm.

